# Impact of occluder device configurations in in-silico left atrial hemodynamics for the analysis of device-related thrombus

**DOI:** 10.1101/2024.01.11.575154

**Authors:** Carlos Albors, Jordi Mill, Andy L. Olivares, Xavier Iriart, Hubert Cochet, Oscar Camara

## Abstract

Left atrial appendage occlusion devices (LAAO) are a feasible alternative for non-valvular atrial fibrillation (AF) patients at high risk of thromboembolic stroke and contraindication to antithrombotic therapies. However, optimal LAAO device configurations (i.e., size, type, location) remain unstandardized due to the large anatomical variability of the left atrial appendage (LAA) morphology, leading to a 4-6% incidence of device-related thrombus (DRT). In-silico simulations have the potential to assess DRT risk and identify the key factors, such as suboptimal device positioning. This work presents fluid simulation results computed on 20 patient-specific left atrial geometries, analysing different commercially available LAAO occluders, including plug-type and pacifier-type devices. In addition, we explored two distinct device positions: 1) the real post-LAAO intervention configuration derived from follow-up imaging; and 2) one covering the pulmonary ridge if it was not achieved during the implantation (13 out of 20). In total, 33 different configurations were analysed. In-silico indices indicating high risk of DRT (e.g., low blood flow velocities and flow complexity around the device) were combined with particle deposition analysis based on a discrete phase model. The obtained results revealed that covering the pulmonary ridge with the LAAO device may be one of the key factors to prevent DRT. Moreover, disk-based devices exhibited enhanced adaptability to various LAA morphologies and, generally, demonstrated a lower risk of abnormal events after LAAO implantation.

## Introduction

Atrial fibrillation (AF) is a life-threatening condition characterized by an irregular electrical conduction and a chaotic contraction. Catalogued as the most prevalent type of arrhythmia, its incidence and prevalence rates steadily rising globally: according to recent statistics, AF affects approximately 33.5 million individuals worldwide with a 20% prevalence in patients aged *>* 80 years [1]. This condition represents a significant clinical challenge due to its associated morbidity and mortality, with an associated substantial socio-economic burden. Among the several associated pathophysiological consequences in AF patients, stroke prevention is a cornerstone of the clinical management of atrial fibrillation, with a reported 5% of stroke risk per year. Structural (i.e., left atrial cavity enlargement) and functional remodeling (i.e., A-wave reduction) produce alterations in blood flow hemodynamics promoting blood stasis and, then, stroke risk. The left atrial appendage (LAA), a remnant ear-shaped and highly trabeculated cavity of the embryonic LA, is the predominant site of thrombosis in AF patients, accounting for over 99% of AF-related strokes [2].

Oral anticoagulants (OACs) related to the K-vitamin antagonist family (i.e., warfarin), or the more recently used Non-vitamin K antagonist (NOACs) oral anticoagulants (i.e., pixaban), are the first-line treatment to reduce and prevent stroke risk. However, an increasing number of non-valvular AF patients have clinical contraindications to (N)OACs, associated with a major risk of bleeding or unexpected side effects due to drug interaction [3]. Over the past decade, left atrial appendage closure using occluder devices, suture loops or excision procedures have emanated as alternatives to prevent and revert the thrombosis process. However, LAA removal or ligation techniques, such as surgical excision and internal suture obliteration, require cardiac arrest, and the use of epicardial clip occlusion or stapled excision can pose challenges related to device navigation and the risk of procedure-related tears along the staple line [4–6].

Several multicentric and randomized clinical trials [7] (EWOLUTION [8], PREVAIL [9], PROTECT-AF [10], ASAP [11], ACP Multicentre [12], Lempereur [13]) have increased the confidence in percutaneous left atrial appendage occlusion (LAAO), reporting non-inferiority to oral anticoagulants. However, successful implantation of a LAAO device remains a challenging endeavor. Among the variability in LAA morphologies and the wide range of device settings to personalize (e.g., design, size, position), the learning curve to acquire excellence in LAAO device implantation becomes extensive, with interventional outcomes being dependent on the operator’s experience [14]. Sub-optimal selection of device characteristics can lead to undesired events at follow-up, such as device embolization, peri-device leaks, or device-related thrombus (DRT). A recent meta-analysis by Alkhouli et al. (2022) [15] reports an incidence of DRT in 4-6% of the cases. The causes leading to DRT may potentially be attributed to various factors, including post-procedural complications (e.g., incomplete device endothelialization), demographic and clinical characteristics (e.g., comorbidities, age-dependency), deep-device-LAA implantation uncovering the pulmonary vein ridge (i.e., *>* 10 mm distance from the Left Superior Pulmonary Vein (LSPV) ridge) or device type (i.e., pacifier- and plug-type devices) [15–17]. While the clinical routine presents the benefits of anticoagulation for clot prophylaxis, there is not a consensus on the ideal drug therapy (i.e., drug choice, dosage, timings) following device implantation, being DRT management still a clinical unsolved problem. In the face of the growing evidence of adverse events, identifying LA morphological and hemodynamics characteristics leading to DRT is key to optimize and simplify LAAO-based therapies.

Over the years, interventional cardiac imaging techniques have been developed to improve the pre-procedural planning and guide left atrial appendage occlusions. The routinely used technique of 2D transesophageal echocardiography (TEE) has been limited by its spatial resolution, being increasingly complemented with more advanced imaging modalities such as 3D TEE and cardiac computed tomography (CCT). While both imaging techniques are currently in use, CCT offers a more comprehensive view of cardiac structures and their spatial relationships, enabling more accurate and reliable anatomical measurements. However, the essential anatomical measurements necessary for selecting the appropriate device, such as the LAA cavity length, width, and diameters and shape of the landing zone and LAA ostium (e.g., interface between the LAA and the main LA cavity), are usually estimated manually from medical images, excessively relying on the operator’s expertise. Moreover, differences in these measurements are observed depending on the selected modality, resulting in variations in device recommendations following manufacturers’ guidelines [18]. Further possibilities for advanced pre-procedural planning are emerging with the use of 3D printing and computational tools such as VIDAA [19], FEOps Heartguide (FEops nv, Gent, Belgium), Mimics (Materialise nv, Leuven, Belgium), WATCHMAN TruPlan (Boston Scientific, Massachusetts, USA), and 3mensio (Pie Medical Imaging bv, Maastricht, Netherlands), which offer direct device setting recommendations based on anatomical estimations. In contrast, these tools do not currently provide insight on blood flow patterns evaluation after the implantation, hindering the prediction and prevention of DRT, as demonstrated in recent joint analysis of morphological and hemodynamics indices [20, 21].

Doppler echocardiography and 4D flow magnetic resonance imaging (MRI) provide certain information about LA hemodynamics such as the quantification of blood flow rates throughout the cardiac cycle. Nonetheless, the former over-simplifies the complex blood flow patterns with a 2D acquisition at a single point. On the other hand, 4D flow MRI has a limited ability to capture rapid changes in blood flow dynamics and identify small or intricate flow features. An emerging alternative approach is the use of cardiac time-resolved CT (4D-CT), which offers better spatio-temporal resolution, but its validation is still in progress [22].

Computational fluid dynamics (CFD) is a powerful tool that can help to fill the gap in understanding LA and LAA morphology and complex hemodynamics characteristics. The hemodynamics of the left atria in patients with atrial fibrillation has been well-studied and -documented in the CFD field, with sensitivity analysis (for a recent 8literature review, see Mill et al. (2021) [23]) establishing best modeling practices for patient-specific simulations [24], analysing rheological conditions [25], input/output boundary conditions, inclusion of wall motion [26–28], among other relevant factors. Nevertheless, despite the increasing number of LAAO interventions, only a limited number of in-silico studies explicitly incorporate LAAO devices in fluid simulations. Some authors have analysed the behavior of blood flow by representing the device surface as a wall [29]. Others have gone further and added preliminary 3D models of the devices [19], including particle adhesion models [30] or residual blood estimations [31]. In parallel, Zaccaria et al. (2020) [32] investigated the mechanical properties of the devices, striving for an optimal in-silico representation of deployment. Even so, these aforementioned studies have been performed in a very limited amount of cases, warranting more comprehensive in-silico studies that include different LAAO device configurations within several patient-specific LA geometries. Such studies are necessary to better understand the relationship with DRT risk. Therefore, the objective of this study was to evaluate the impact in the LA hemodynamics of the device type (i.e., pacifier versus plug shape) and location (i.e., coverage versus non-coverage of pulmonary ridge), and their potential relation with device-related thrombus formation in 20 patient-specific left atrial geometries with pre-procedural and follow-up CT scans of patients who underwent a LAAO device implantation. Computer-aided design models of pacifier- and plug-type devices were virtually implanted, simulating the implanted device configuration from the follow-up CT scans. Subsequently, in those cases where the pulmonary ridge (PR) was not fully covered with the implanted device, we simulated an alternative LAAO configuration to analyse the influence of PR coverage on DRT risk.

## Materials and methods

### Clinical data

Retrospective pre- and post-occlusion computed tomography images of 20 non-valvular AF patients were provided by Hôpital Haut-Lévêque (Bordeaux, France), after approval from the ethical committee and informed consent of the patients. Cardiac CT studies acquired on a 64-slice dual-source CT system (Siemens Definition, Siemens Medical Systems, Forchheim, Germany) and then reconstructed into isotropic voxel sizes (0.37-0.5 mm range; 512 × 512 × [270-403] slices). The six-month follow-up CT scan showed the presence of a device-related thrombus in 10 of the 20 patients, following the degree of hypoattenuating thickening (HAT) in the classification detailed in Cochet et al. (2018) [33] study (see Table 1).

**Table 1.** Characterization of the cohort of 20 patients analysed in terms of clinical outcomes related to control and device-related thrombosis (DRT) subjects, with the device type and size. The analysis covers the implanted post-left atrial appendage occlusion (LAAO) configuration, as well as the proposed configuration covering the pulmonary ridge (PR). Device types were maintained on the configuration covering the pulmonary ridge. ACP: Amplatzer Cardiac Plug.

|  | Clinical post-LAAO outcome | Post-LAAO configuration |  | PR covered configuration | LA volume (ml) |
| --- | --- | --- | --- | --- | --- |
|  |  | Device type | Device size | Device size |  |
| Patient #1 | Control | Amulet | 25 mm | 28 mm | 171 |
| Patient #4 | Control | Watchman | 33 mm | 27 mm | 181 |
| Patient #5 | Control | Watchman | 27 mm | - | 216 |
| Patient #6 | Control | Watchman | 27 mm | - | 94 |
| Patient #7 | Control | Amulet | 25 mm | - | 179 |
| Patient #8 | Control | Amulet | 28 mm | - | 196 |
| Patient #9 | Control | Watchman | 24 mm | 30 mm | 158 |
| Patient #11 | Control | Amulet | 22 mm | - | 127 |
| Patient #13 | Control | Watchman | 24 mm | 24 mm | 214 |
| Patient #20 | Control | Watchman | 24 mm | 27 mm | 100 |
| Patient #2* | DRT | ACP | 22 mm | 28 mm | 161 |
| Patient #3* | DRT | ACP | 16 mm | 20 mm | 109 |
| Patient #10* | DRT | Watchman | 24 mm | 27 mm | 89 |
| Patient #12* | DRT | Amulet | 25 mm | - | 212 |
| Patient #14* | DRT | Amulet | 25 mm | 34 mm | 242 |
| Patient #15* | DRT | Watchman | 30 mm | 33 mm | 206 |
| Patient #16* | DRT | Watchman | 24 mm | 27 mm | 225 |
| Patient #17* | DRT | ACP | 28 mm | 26 mm | 294 |
| Patient #18* | DRT | ACP | 28 mm | - | 213 |
| Patient #19* | DRT | Watchman | 27 mm | 30 mm | 220 |

### Occluder deployment process and patient-specific cohort generation

Binary masks of the left atrial geometry were segmented from the CT images using semi-automatic threshold tools available in Slicer 4.10.11 (https://www.slicer.org/). Subsequently, pre- and post-occlusion patient-specific LA surface meshes were reconstructed with the flying edges algorithm also in Slicer. In addition, the implanted device was manually segmented from the post-CT images, representing the real LAAO configuration at follow-up for each analysed case. As the characterization of the LAA geometry cannot be extracted from post-occlusion CT images (due to lack of contrast), the reconstructed implanted device was registered to the pre-occlusion LA geometry to replicate in-silico the LAAO configuration. To do so, a fiducial rigid registration from Meshlab v2021-07 (https://www.meshlab.net/) between both pre- and post-LAAO meshes were performed, using an average of 8 manually selected landmarks in the pulmonary veins (PV) antrum, LAA ostium, and, mitral valve (MV) annulus. The rigid registration method was based only on rotations and transformations without changes of scale.

Then, the web-based VIDAA platform [19] was used to place the device in the pre-LAAO mesh, building computer-aided design (CAD) occluder models from the segmented device as a reference. The device segmentation, in conjunction with clinical records, facilitated the identification of the device size, type, and location from the post-CT images, which were then used to select the appropriate CAD model of the device to deploy in the pre-CT LA geometry. The device was manually deployed after the VIDAA platform’s recommendations for landing zone and device size. Each final device configuration was reviewed by an interventional cardiologist to ensure a plausible deployment. The cohort of the present study employed three frequently used occluders to evaluate the performance of device type: the Amplatzer Amulet and Cardiac Plug (St. Jude Medical-Abbott, St. Paul, Minnesota, United States), which are both pacifier-type devices, and the Watchman (Boston Scientific, Marlborough, Massachusetts, United States), which is a plug-type device. CAD models of LAAO devices were necessary due to the limited spatial resolution of the CT scan, which prevented a detailed reconstruction of the implanted device, especially in the plug device where only the metallic structure was visible (see Fig. 2). Moreover, the hangs of the plug-type device were removed after device deployment to reduce the computational cost of the simulation.

For some patients, a second LAAO device configuration (see plug-type device in Fig. 1), different from the implanted one, was developed to better cover the pulmonary ridge (lateral fold results from the merging of the LAA and LSPV) in the LAA [34] and analyse potential variations in blood flow patterns. To determine the appropriate size for this configuration, the VIDAA platform provided recommendations based on anatomical measurements, in accordance with the device manufacturers’ guidelines [18].

**Fig 1.**
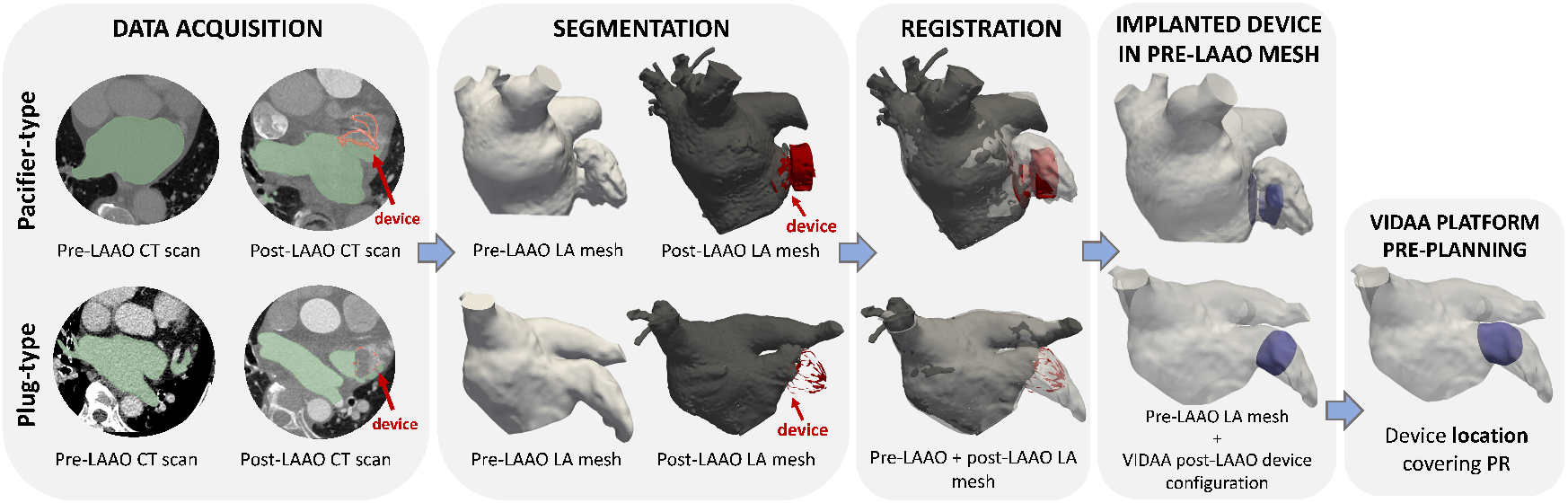
Modeling pipeline of plug-type and pacifier-type device deployment in the pre-operative left atrial geometry. A proposed configuration covering the pulmonary ridge (PR) is suggested in the plug-type device. LAAO: Left atrial appendage occlusion; CT: computed tomography. VIDAA: Virtual Implantation and Device selection in left Atrial Appendages.

**Fig 2.**
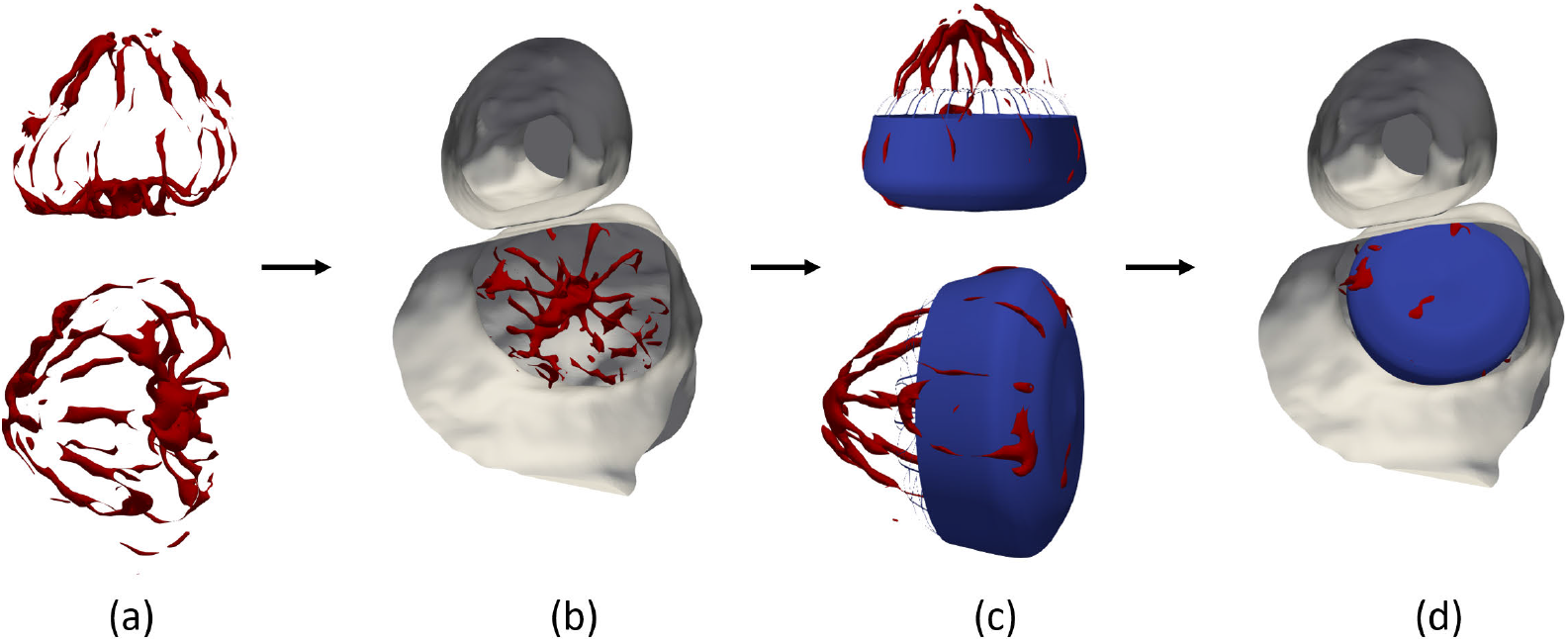
Device deployment process of a plug-type device. (a) Segmented plug-type device (red wireframe) from computed tomography (CT) scans following a left atrial appendage occlusion (LAAO) procedure. (b) LAAO device configuration in a longitudinal LAA cut of the CT scan. (c) Computer-aided design model (blue) of the plug-type device replicating the LAAO configuration extracted from the CT scan. (d) Final device configuration within the LAA cavity.

Post-processing steps were applied using Meshlab to the pre-LAAO LA surface meshes and CAD models to correct intersecting faces and no-manifold edges out of the segmentation and device deployment process. Also, planar surfaces at the end of the pulmonary veins and mitral valve were defined in Autodesk Meshmixer v3.3.15 (https://meshmixer.com/), so the simulated blood flow definition was simpler and perpendicular to this surface (normal vector). In total, 20 pre-occlusion surface left atrial meshes defining 33 different device configurations were composed of*≈* 7 *×* 10^4^ surface elements with an average edge length of 0.68 mm (see Table 1 for more details on configurations).

Finally, a 3D Delaunay refinement algorithm with a Netgen [35] tetrahedra quality optimization from Gmsh 4.5.4 software (https://gmsh.info/) were employed to construct the volumetric meshes to solve the fluid domain. The tetrahedral volumetric mesh resolution (1, 2 *×* 10^6^ elements) was determined following recent sensitivity studies [20, 23, 24] in the field.

## In-silico simulations

### Boundary conditions and setup of simulations

Computational fluid dynamic simulations were performed using the Ansys Fluent 2022 solver (ANSYS Inc, USA). Blood rheology was considered as an incompressible Newtonian fluid with a density of *ρ* = 1060 kg m^*−*3^ and viscosity of *μ* = 0.0035 kg/m*s in a laminar regime [36]. Generic clinical measurements collected from an AF patient used as boundary conditions were equally in all the cases, based on previous analysis on LA-based fluid models [23]. A catheter pressure data profile extracted in sinus rhythm was applied at the pulmonary veins (i.e., inlets), and velocity dynamics derived from Doppler echocardiography was imposed at the mitral valve (i.e., outlet). The simulation was run for two cardiac cycles, with 88 steps per beat and a Δ*t* = 0.01*s*, in accordance with the patient’s heart rate (HR). Moreover, a half-scaled function representing the mitral valve annulus displacement, from Veronesi et al. [37], with a peak displacement of 4 mm was set as in [28]. Then, LA wall motion diffusion was estimated with a spring-based dynamic mesh approach within the CFD solver [23]. The average computational time required for each simulation was 25 h with an Intel i9-9900k CPU and 32GB RAM desktop performed in serial.

### Platelet adhesion estimation using a discrete phase modeling

The process of thrombus formation was approximated based on a discrete phase model (DPM) coupled with the CFD solver, simulating platelet adhesion which is one of the initial steps in the molecular mechanism of thrombus formation. The blood flow continuous phase (Eulerian approach) was combined with the discrete phase (Lagrangian approach) through the injection of a given number of particles. The interaction was represented by tracking a number of particles injected through the pulmonary veins and dragged by the flow until their attachment to the endothelial wall of the LAA or their exit through the MV. Moreover, momentum and energy could also be exchanged between the Eulerian and Lagrangian approaches. Individual particle trajectories, *p*, were computed with the integration of the force balance within the continuous phase. This can be expressed in the general form as:

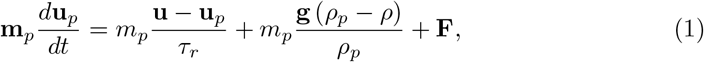

where **u** describes fluid domain velocity; **m**_*p*_, *ρ*_*p*_, and **u**_*p*_ corresponds to the particle mass, density and velocity, respectively; *τ*_*r*_ is the droplet relaxation time, and *ρ* is the blood density. In Equation 1, different forces have been considered: the drag force (*m*_*p*_ (**u** *−* **u**_*p*_) */τ*_*r*_), pressure gradient forces, *F*_*pg*_, and virtual mass forces *F*_*vm*_ to accelerate the fluid enclosing the particles, defined as:

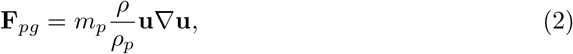

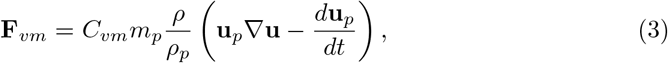

where the virtual mass factor, *C*_*vm*_, is defined as 0.5. Gravitational forces were neglected.

Moreover, a Saffman lift force [38] term was added within the DPM to represent the lift due to shear. Shear lift [39] is caused by inertial effects in viscous flows around the particle and is defined as follows:

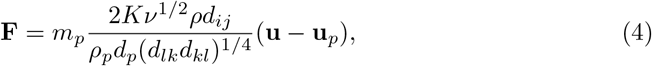

where *d*_*ij*_ as the deformation tensor, *ν* the kinematic viscosity and the constant coefficient *K* =2.954 [39].

### Particle injections

A pre-defined number of particles per case representing a cluster of platelets were injected through the planar surfaces of the PVs during the initial 10 time steps of each simulated cardiac cycle beat. Physiological conditions for blood platelet concentration, *c*_*p*_, has been assumed ^1^; hence a *c*_*p*_ value of 2 10^8^ mL^*−*1^ were imposed during the initial 10 time steps at the beginning of the simulation. Then, to guarantee the physiological platelet concentration, parameters like platelets per cluster, *n*_*ppc*_, total flow rate, *Q*_*flow*_, and particle diameter, *d*_*p*_, were personalized in each analysed case, based on anatomical characteristics. For instance, the number of platelets per cluster, *n*_*ppc*_, was computed as:

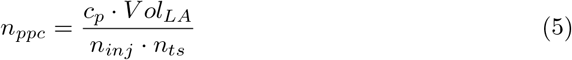

where *V ol*_*LA*_ is the LA volume, *n*_*ij*_ the number of facets of the inlet planar PV surfaces, and *n*_*ts*_, the number of injection time steps. The enumerator defines the number of platelets per injection while the denominator represents the number of platelets injected. Assuming spherical shape on the clusters and platelets, the platelet diameter, *d*_*p*_, can be estimated as follows:

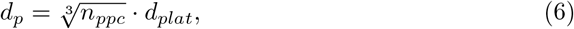

where Equation 6 is composed of the platelet diameter *d*_*plat*_ extracted from the particle volume equation 7:

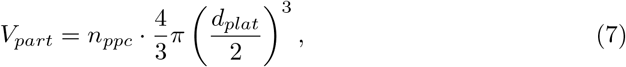

and with the definition of the particle diameter *d*_*p*_ as:

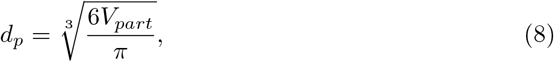

Finally, the total flow rate, *Q*, was described by the mass injected per unit of time as follows:

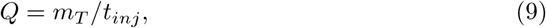

where *m*_*T*_ is the total mass added within the whole injection:

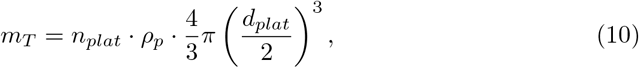

and *t*_*inj*_ is the injection time:

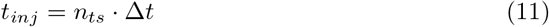

The previous equations were solved by assuming the pre-determinated values [40]: *n*_*ts*_ = 10, *d*_*plat*_ = 3 *μ*m, particle density *ρ*_*p*_ = 1550 kg m^*−*3^, and a time step size of Δ*t* = 0.01 s. In addition, a particle velocity of 0.01 m/s was specified for each injection step to ensure sufficient inertia for the particles to be carried by the fluid flow. Identical parameters were imposed at the initial steps of the second injection in the second cardiac cycle.

### Particle attachment

A worst-case scenario for DRT was assumed through the platelet adhesion model, in which all particles touching the endothelial LAA wall were adhered by the wall-film constraint, independently of the velocity magnitude. No splashing of particles was assumed, so the shape was maintained. However, the model considered an eventual detachment scenario with the O’Rourke [41] separation model or a roll-over of the particles over the LAA surface determined by the platelets properties of surface tension *σ*_*p*_ = 0.03 N/m and molecular viscosity *μ*_*p*_ = 0.0025 kg m^*−*1^ s^*−*1^. The estimation of the thrombus formation risk was based on the evaluation of the platelets’ number and position.

### Hemodynamic indices

To complement the DRT evaluation, a range of in-silico indices were derived from simulation results to characterize the hemodynamic differences across the 33 device configurations. Specifically, the measurement area encompassed the region from the fold of the left superior pulmonary vein to the device surface, which constitutes a susceptible zone for low flow velocities (i.e., *<* 0.2 m/s) and complex fluid dynamics [42].

Qualitative and quantitative evaluation of the simulated blood flow pattern was conducted via streamline representation and through the computation of the averaged velocity in the targeted region around the device, respectively. Moreover, the endothelial cell activation potential (ECAP) index [43] was calculated to detect endothelial injury, a precursor to inflammation and thrombus formation. The computation of ECAP involved determining the ratio between the time-averaged wall shear stress (TAWSS) associated with flow velocities and the oscillatory shear index (OSI) associated with flow complexity. The OSI values ranged from 0, indicating unidirectional flow, to 0.5, indicating bidirectional flow. ECAP values were anticipated to be higher in regions with low blood flow velocities and complex fluid patterns, indicating a greater risk of thrombus formation. The analysis was performed during key cardiac cycle phases (late-systole and early-, and late-diastole) in the second cardiac beat to minimize convergence issues. The resulting simulation data were post-processed, visualized, and analysed using ParaView version 5.7.0 ^2^.

## Results

### Device configuration results

Table 1 presents the 20 configurations extracted from post-LAAO CT images. The cohort was evenly distributed between patients with plug- and pacifier-type occluders, with 10 patients in each group. The control cases were composed of 6 patients with a plug-type device and 4 patients with a pacifier-type device, while the DRT group were distributed in the opposite manner.

Thirteen additional device configurations were simulated to cover the pulmonary ridge in those cases where the implanted device did not cover it well (i.e., deep device implantations of the original configuration or possible better device surface alignment with the ostium plane). Out of the 13 cases, 8 corresponded to patients having DRT after LAAO implantation. The same device type as the original configuration was used in each proposed configuration. Except for patients #4 and #17*, larger device sizes were virtually implanted following manufacturers’ recommendations for increased anatomical dimensions of the landing zone. In addition, the 20 analysed patients encompass the large variability of the left atrial appendage morphologies, including cases from all categories in the classical classification proposed by Di Biase et al. (2012) [44]. Moreover, there was heterogeneity in pulmonary vein configuration (i.e., 14 patients with four PVs, three patients with five PVs, and three patients with six PVs) which may potentially affect blood behavior patterns, as recently suggested [45]. There were no statistically significant distinctions in left atrial volumes between the control (163.6 ± 39 ml) and DRT group (177.1 ± 58 ml) relating larger LA remodeling with a higher risk of DRT.

### Simulated blood flow patterns

A preliminary analysis assessed the influence of the DPM inclusion on blood flow behavior in computational simulations, revealing minor changes in flow patterns without significant impact on the interpretation or estimation of DRT. The interested reader is referred to Appendix:Discrete phase model effects on blood flow behaviour for further information.

In the real post-LAAO configuration, comparable averaged blood flow velocity were observed within both study groups, encompassing all cases within each respective group (0.2 *±* 0.11 for the control group and 0.195 *±* 0.12 for the DRT group). The simulated blood flow (see Table 2, Fig. 3, and Supplementary material Supplementary Material) revealed that 12 patients (patients #1, #2*, #3*, #5, #6, #9, #10*, #13, #15*, #16*, #19*, and #20) exhibited re-circulation patterns with velocities below the threshold value of 0.2 m/s on the device surface at several instants of the cardiac cycle. This threshold is typically associated with a high risk of blood stasis [42]. These patients were equally distributed between the control and DRT groups. However, in two of the control cases (patients #5, and #6) and one DRT case (Patient #10*), good washout was observed in the high-risk regions of thrombosis where high flow velocities (*>* 0.2 m/s) with laminar behavior were maintained for a significant portion of the mid-, late-diastolic phase. The remaining nine cases (patients #1, #2*, #3*, #9, #13, #15*, #16*, #19*, and #20) depict flow velocities below the risk value of 0.2 m/s when considering the average of the entire cardiac cycle. Among this subgroup with low average velocities (5 DRT and 4 control cases), the predominant characteristic was the presence of an uncovered pulmonary ridge, except for patient #16*. In contrast, the group of patients with mean flow velocities above 0.2 m/s (patients #4, #5, #6, #7, #8, #10*, #11, #12*, #14*, #17*, and #18*), only three (patients #4, #10*, and #14*) had an uncovered pulmonary ridge. Moreover, two peri-device leaks were identified in patients #2* and #7.

**Table 2.**
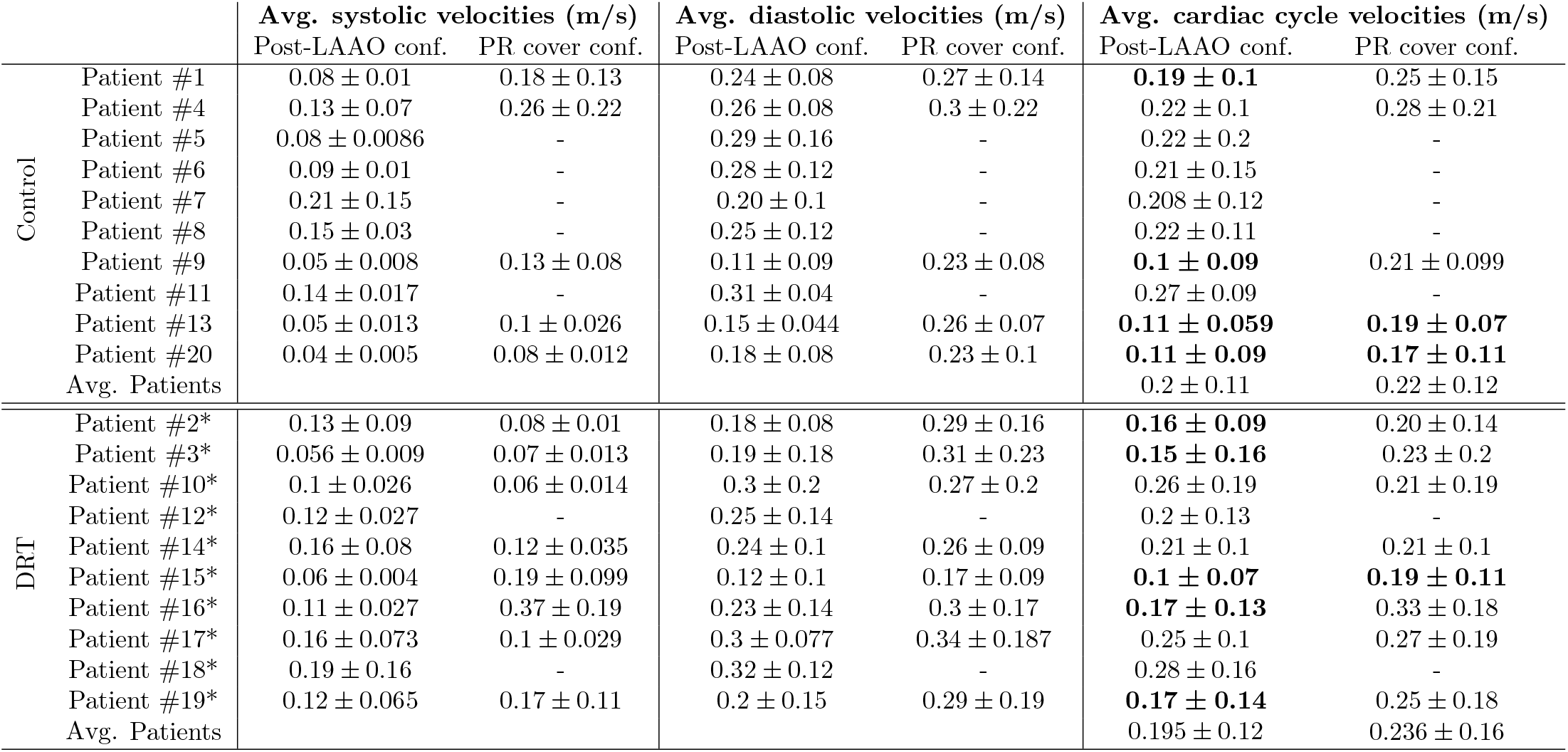
Averaged (Avg.) velocities (average *±* standard deviation) during systolic, diastolic and full cardiac cycle phases on device surface in the cohort of 20 cases. Mean velocity values of the entire cardiac cycle *<* 0.2 m/s (highlighted in bold) may indicate high risk of device-related thrombosis (DRT) [42]. LAAO: left atrial appendage occlusion. PR: pulmonary ridge. * defines the patients with DRT.

|  |  | Avg. systolic velocities (m/s) |  | Avg. diastolic velocities (m/s) |  | Avg. cardiac cycle velocities (m/s) |  |
| --- | --- | --- | --- | --- | --- | --- | --- |
|  |  | Post-LAAO conf. | PR cover conf. | Post-LAAO conf. | PR cover conf. | Post-LAAO conf. | PR cover conf. |
| Control | Patient #1 | $0.08 \pm 0.01$ | $0.18 \pm 0.13$ | $0.24 \pm 0.08$ | $0.27 \pm 0.14$ | **$0.19 \pm 0.1$** | $0.25 \pm 0.15$ |
| | Patient #4 | $0.13 \pm 0.07$ | $0.26 \pm 0.22$ | $0.26 \pm 0.08$ | $0.3 \pm 0.22$ | $0.22 \pm 0.1$ | $0.28 \pm 0.21$ |
| | Patient #5 | $0.08 \pm 0.0086$ | - | $0.29 \pm 0.16$ | - | $0.22 \pm 0.2$ | - |
| | Patient #6 | $0.09 \pm 0.01$ | - | $0.28 \pm 0.12$ | - | $0.21 \pm 0.15$ | - |
| | Patient #7 | $0.21 \pm 0.15$ | - | $0.20 \pm 0.1$ | - | $0.208 \pm 0.12$ | - |
| | Patient #8 | $0.15 \pm 0.03$ | - | $0.25 \pm 0.12$ | - | $0.22 \pm 0.11$ | - |
| | Patient #9 | $0.05 \pm 0.008$ | $0.13 \pm 0.08$ | $0.11 \pm 0.09$ | $0.23 \pm 0.08$ | **$0.1 \pm 0.09$** | $0.21 \pm 0.099$ |
| | Patient #11 | $0.14 \pm 0.017$ | - | $0.31 \pm 0.04$ | - | $0.27 \pm 0.09$ | - |
| | Patient #13 | $0.05 \pm 0.013$ | $0.1 \pm 0.026$ | $0.15 \pm 0.044$ | $0.26 \pm 0.07$ | **$0.11 \pm 0.059$** | **$0.19 \pm 0.07$** |
| | Patient #20 | $0.04 \pm 0.005$ | $0.08 \pm 0.012$ | $0.18 \pm 0.08$ | $0.23 \pm 0.1$ | **$0.11 \pm 0.09$** | **$0.17 \pm 0.11$** |
| Avg. Patients | | | | | | $0.2 \pm 0.11$ | $0.22 \pm 0.12$ |
| DRT | Patient #2* | $0.13 \pm 0.09$ | $0.08 \pm 0.01$ | $0.18 \pm 0.08$ | $0.29 \pm 0.16$ | **$0.16 \pm 0.09$** | $0.20 \pm 0.14$ |
| | Patient #3* | $0.056 \pm 0.009$ | $0.07 \pm 0.013$ | $0.19 \pm 0.18$ | $0.31 \pm 0.23$ | **$0.15 \pm 0.16$** | $0.23 \pm 0.2$ |
| | Patient #10* | $0.1 \pm 0.026$ | $0.06 \pm 0.014$ | $0.3 \pm 0.2$ | $0.27 \pm 0.2$ | $0.26 \pm 0.19$ | $0.21 \pm 0.19$ |
| | Patient #12* | $0.12 \pm 0.027$ | - | $0.25 \pm 0.14$ | - | $0.2 \pm 0.13$ | - |
| | Patient #14* | $0.16 \pm 0.08$ | $0.12 \pm 0.035$ | $0.24 \pm 0.1$ | $0.26 \pm 0.09$ | $0.21 \pm 0.1$ | $0.21 \pm 0.1$ |
| | Patient #15* | $0.06 \pm 0.004$ | $0.19 \pm 0.099$ | $0.12 \pm 0.1$ | $0.17 \pm 0.09$ | **$0.1 \pm 0.07$** | **$0.19 \pm 0.11$** |
| | Patient #16* | $0.11 \pm 0.027$ | $0.37 \pm 0.19$ | $0.23 \pm 0.14$ | $0.3 \pm 0.17$ | **$0.17 \pm 0.13$** | $0.33 \pm 0.18$ |
| | Patient #17* | $0.16 \pm 0.073$ | $0.1 \pm 0.029$ | $0.3 \pm 0.077$ | $0.34 \pm 0.187$ | $0.25 \pm 0.1$ | $0.27 \pm 0.19$ |
| | Patient #18* | $0.19 \pm 0.16$ | - | $0.32 \pm 0.12$ | - | $0.28 \pm 0.16$ | - |
| | Patient #19* | $0.12 \pm 0.065$ | $0.17 \pm 0.11$ | $0.2 \pm 0.15$ | $0.29 \pm 0.19$ | **$0.17 \pm 0.14$** | $0.25 \pm 0.18$ |
| Avg. Patients | | | | | | $0.195 \pm 0.12$ | $0.236 \pm 0.16$ |

**Fig 3.**
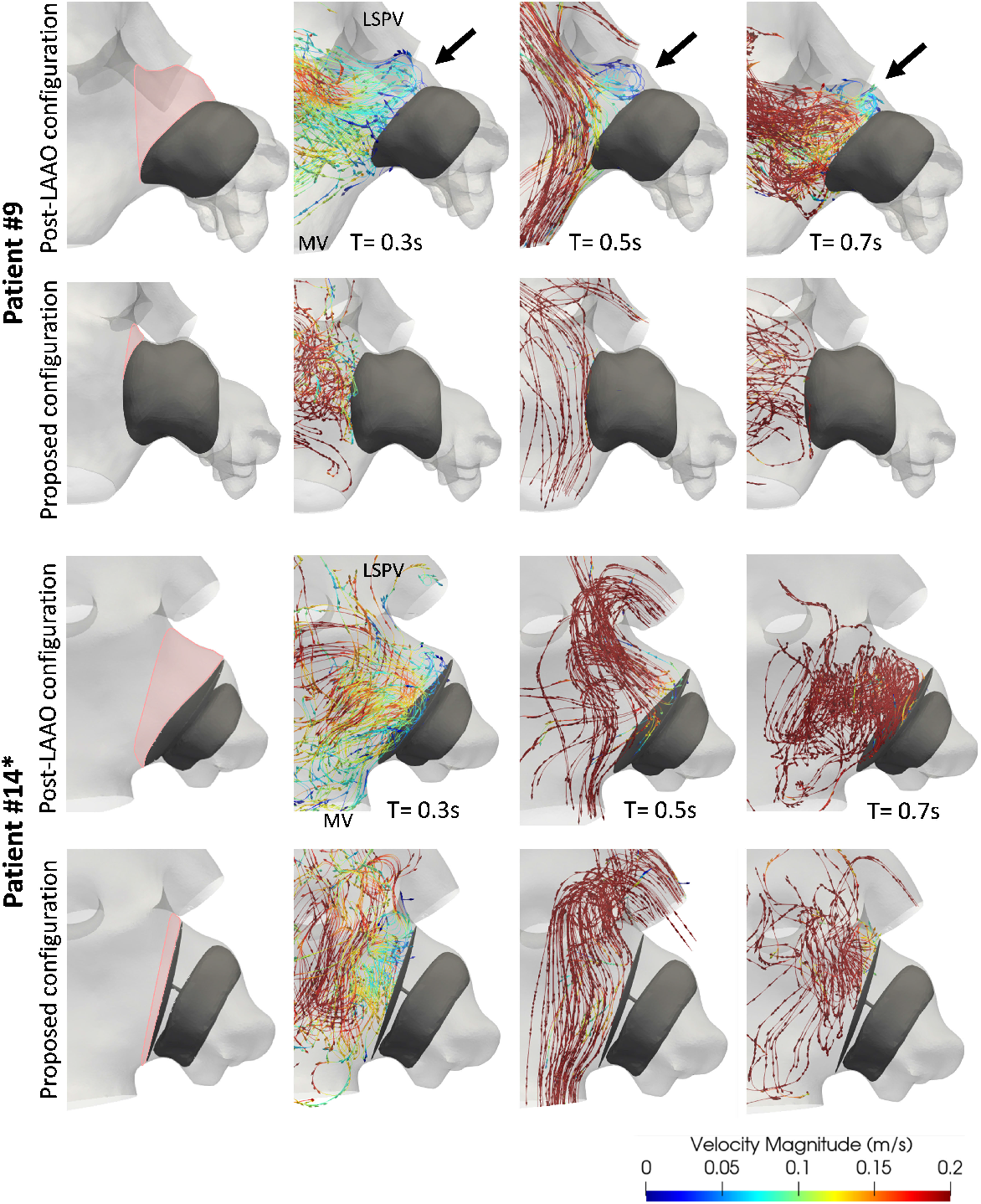
Simulated fluid flow patterns during late systole (t = 0.3 s), early diastole (t = 0.5 s), and late diastole (t = 0.7 s) in the control patient #9 and patient #14* with device-related thrombus (DRT). The first column depicts the uncovered region of the pulmonary ridge in pink. LAAO: left atrial appendage occlusion. LSPV: left superior pulmonary vein. MV: mitral valve. * indicates patients with DRT.

In patients where a proximal configuration covering the pulmonary ridge was simulated, there was a significant reduction in re-circulation flows observed in the majority of cases (see Table 2, Fig. 3, and Supplementary material Supplementary Material). Additionally, average velocities were found to be higher in the proposed position in comparison to the original one in all cases. However, for patients #13, #15*, and #20, average velocities higher than the threshold of 0.2 m/s were not achieved. In general, during the diastolic phase, higher velocities were observed regardless of the device configuration, with a greater amount of re-circulations at low velocities during the systolic phase.

When analysing the device types, the plug-type occluder device was found to be predominant in the patients with low average velocities and complex patterns on the device surface of the real post-LAAO configuration, accounting for 9 out of the 12 patients (patients #5, #6, #9, #10*,#13, #15*, #16*, #19*, and #20). This trend was particularly observed in the proposed device configuration covering the pulmonary ridge, where the 3 cases with lower average velocities had the plug-type device implanted.

For a more detailed analysis, Figure 3 displays the control Patient #9 and DRT Patient #14*, both exhibiting an uncovered pulmonary ridge. The control case provides a clear example of low velocity re-circulations on the surface of the device with an uncovered pulmonary ridge. Complex patterns were observed on the device in all three instants (late-systole, early- and late diastole) of the cardiac cycle. However, the patterns transformed into a more laminar behavior and higher velocities with a covered pulmonary ridge. In contrast, Patient #14* only showed low blood flow velocities and re-circulations during the systolic phase (t = 0 - 0.3 s), which were replaced by high velocity flows during the entire diastolic phase (t = 0.31 s - 0.88 s). When covering the pulmonary ridge, the behavior remained the same, and even lower velocities were detected on device disk at the end of the cycle (t = 0.7 s).

## Endothelial cell activation potential (ECAP)

The in-silico index estimating thrombogenic risk, endothelial cell activation potential, was evaluated across the 33 device configurations, allowing us to identify regions with a high thrombogenic risk (*>* 0.5 Pa^*−*1^ [43, 46]) within the study cohort. Fourteen patients had maximum ECAP values *>* 0.5 Pa^*−*1^ in the original post-LAAO implantation, with the control group standing out with 8 patients (see Table 3 Post-LAAO config.). In addition, the control group exhibited higher peak ECAP values compared to the DRT group above the ECAP threshold value analysed (1.25 *±* 0.41 for the control group versus 0.98 *±* 0.18 for the DRT group), due to high OSI values. Notably, only patients #1, #14*, and #19 showed small, localized regions with high ECAP values, while the remaining patients had them distributed over the entire area adjacent to the occluder device (see Fig. A.2). patients #2*, #4, #13, and #20 stood out for having a mean value close to or even exceeding the threshold value. Similarly to the velocity findings, 10 out of the 14 patients with high ECAP values did not have a covered PR and a plug-type occluder device was implanted, especially in the control group where 6 of the 8 patients had the plug-type device.

**Table 3.** Peak and mean (average *±* standard deviation) values of the endothelial cell activation potential (ECAP) on device surface in the cohort of 20 cases. ECAP values *>* 0.5 Pa^*−*1^ (highlighted in bold) may indicate high risk of device-related thrombus (DRT). LAAO: left atrial appendage occlusion. PR: pulmonary ridge * defines the patients with DRT.

| | | Max. ECAP value ($\text{Pa}^{-1}$) | | Mean ECAP value ($\text{Pa}^{-1}$) | |
| --- | --- | --- | --- | --- | --- |
|  |  | Post-LAAO conf. | PR cover conf. | Post-LAAO conf. | PR cover conf. |
| Control | Patient #1 | 0.68 | 0.51 | $0.24 \pm 0.19$ | $0.2 \pm 0.23$ |
| | Patient #4 | 1.24 | 0.44 | $0.46 \pm 0.9$ | $0.11 \pm 0.08$ |
| | Patient #5 | 0.7 | - | $0.26 \pm 0.17$ | - |
|  | Patient #6 | 1.55 | - | **$0.6 \pm 0.7$** | - |
| | Patient #7 | 0.39 | - | $0.19 \pm 0.08$ | - |
| | Patient #8 | 0.43 | - | $0.14 \pm 0.1$ | - |
| | Patient #9 | 1.42 | 0.63 | $0.3 \pm 0.2$ | $0.24 \pm 0.13$ |
| | Patient #11 | 1.02 | - | $0.3 \pm 0.4$ | - |
| | Patient #13 | 1.4 | 1.2 | **$0.7 \pm 0.3$** | $0.45 \pm 0.43$ |
|  | Patient #20 | 2 | 1.4 | **$1.1 \pm 1.3$** | **$0.6 \pm 0.52$** |
| DRT | Patient #2* | 0.99 | 0.48 | **$0.49 \pm 0.4$** | $0.19 \pm 0.1$ |
| | Patient #3* | 0.55 | 0.22 | $0.19 \pm 0.13$ | $0.14 \pm 0.143$ |
| | Patient #10* | 1.26 | 0.55 | $0.23 \pm 0.2$ | $0.11 \pm 0.14$ |
| | Patient #12* | 0.48 | - | $0.08 \pm 0.1$ | - |
| | Patient #14* | 0.8 | 0.3 | $0.32 \pm 0.12$ | $0.19 \pm 0.2$ |
| | Patient #15* | 0.76 | 0.3 | $0.33 \pm 0.2$ | $0.26 \pm 0.4$ |
| | Patient #16* | 1.08 | 0.51 | **$0.8 \pm 0.6$** | $0.13 \pm 0.2$ |
| | Patient #17* | 0.3 | 0.32 | $0.14 \pm 0.11$ | $0.15 \pm 0.087$ |
| | Patient #18* | 0.2 | - | $0.16 \pm 21$ | - |
| | Patient #19* | 0.73 | 0.6 | $0.24 \pm 0.33$ | $0.29 \pm 0.2$ |

Positioning the device covering the pulmonary ridge (see Table 3 PR cover config.) reduced the number of patients with maximum ECAP values above 0.5 Pa^*−*1^ from 14 to four (patients #9, #13, #19*, and #20). While the ECAP values were generally reduced compared to the original positions, broad and well-defined regions with ECAP values exceeding 1 were still observed in patients #13 and #20 (see Fig. A.2).

## Assessment of particle models

The inclusion of the DPM with the fluid domain solution facilitates a preliminary estimation of the coagulability risk associated with the visualization of attachment-prone regions, platelet quantity, and attachment velocity within the left atrial appendage. The average attachment velocities (see Fig. A.3) showed a wide range of velocities before the particles became stuck to the wall. Generally, the highest accumulation occurred in mid-diastole, coinciding with the maximum blood velocity driven by the E-wave, which is attributed to the increased influx of platelets carried by the fluid within the LAA.

The control cases, in comparison to the DRT group, demonstrate a lower percentage of adhesion (see Fig. 4). Notably, patients #10* and #15* within the DRT group exhibit the highest peak of adhesion. The mean percentage of adhesion in the control group was 4.8%, whereas, in the DRT group, it was 11.4%. In terms of device configurations, as shown in Figure 4, the model revealed greater platelet accumulation in patients with an uncovered pulmonary ridge ostium, particularly in patients #1, #2*, #10*, #14*,#15*, #17* and, #20, who exhibited more than 10% platelet accumulation at the same instant of time (late diastole T = 0.88 s). Patient 16* was the only one who did not show improvement with the proposed position covering the pulmonary ridge.

**Fig 4.**
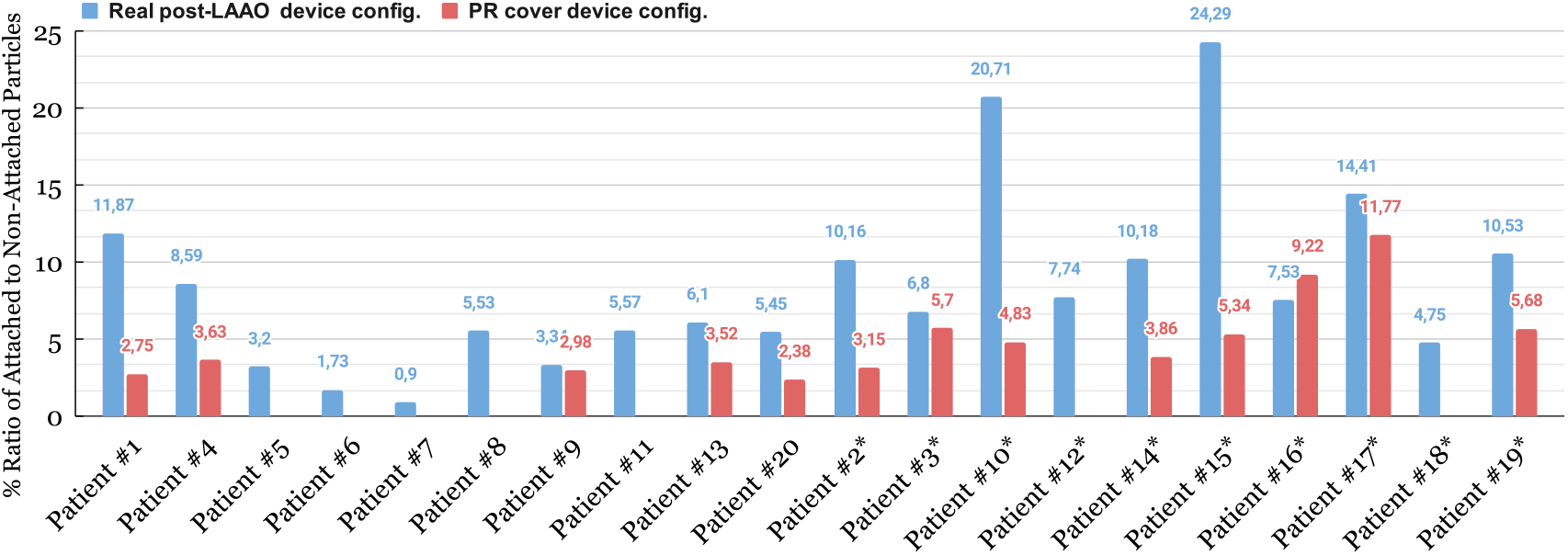
Quantification of particle attachment to the left atrial appendage wall at the end of the cardiac cycle in the 20 patients cohort. Blue represents the post-left atrial appendage occlusion (LAAO) configuration and, red represents the proposed configuration covering the pulmonary ridge. PR: pulmonary ridge. * denotes patients with device-related thrombus (DRT).

In the original post-LAAO configuration, the velocity of the particles prior to adhesion to the wall was *<* 0.4 m/s, except in patients #18* and #10*, which were higher. In the proposed location covering the PR, the velocities before the attachment were higher due to the device’s position and the presence of laminar flows on the device surface. Once the particles contacted the wall, regardless of the position, the velocity decreased significantly (*<* 0.06 m/s) due to the imposed wall-film condition. The effect of detachment and drag on the wall was observed in cases where the PR was covered, as previously adhered particles showed an increase in velocity (up to 0.2 m/s) while rolling through the wall. Two peri-device leaks were detected from the simulated flows in the original post-LAAO configuration: Patient #2* having a major leak with significant platelet accumulation in the distal lobes of the LAA (see Fig. 5 Patient #2* (a)); and Patient #7 having a minor leak with low platelets behind the device.

**Fig 5.**
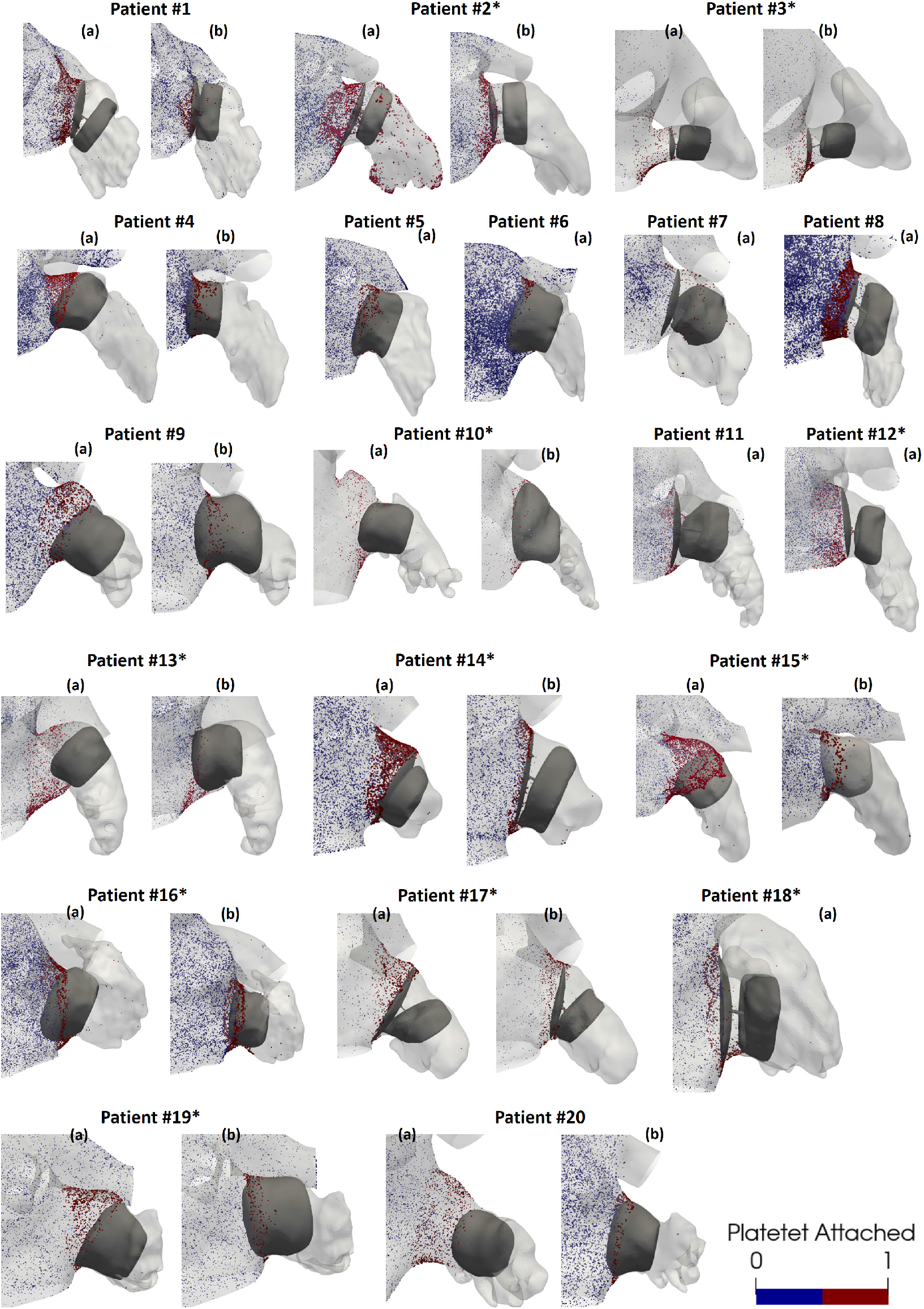
The Discrete Phase Model (DPM) results for the 33 analysed configurations, showing platelet concentration from a frontal visualization of the left atrial appendage (LAA). Panel (a) represents the real post-left atrial appendage occlusion (LAAO) configuration, while panel (b) shows the proposed configuration covering the pulmonary ridge. * defines the patients with DRT

Although platelets were observed in the vicinity of both device types (see Fig. 5), the plug-type device exhibited more platelet accumulation in certain regions, including the curvature zone between the device’s lobe and the endothelial wall of the LAA. These effects were evident even in cases where the PR was covered, as in patients #5, #6, and #16*. The pacifier device showed a homogeneous distribution of platelets on its entire surface (i.e., patients #2*, #3*, #14*) with patient #1 being the exception in the proposed location with the PR covered.

## Overview results

The synthesised findings of Table 4 revealed that only in nine patients, the hemodynamic descriptors and particle assessment successfully captured the clinical outcome. Among them, five patients (#2*, #7, #8, #15*, and #19*) exhibited complete agreement between all four indices and the clinical outcome. The other four patients (#4, #10*, #11, and #16*) demonstrated agreement with three out of the four descriptors and the clinical outcome. Surprisingly, in patients #12* and #18* from the DRT group, none of the studied descriptors provided an estimation of high risk for DRT. The DPM yielded the most favorable results out of all the descriptors evaluated.

**Table 4.**
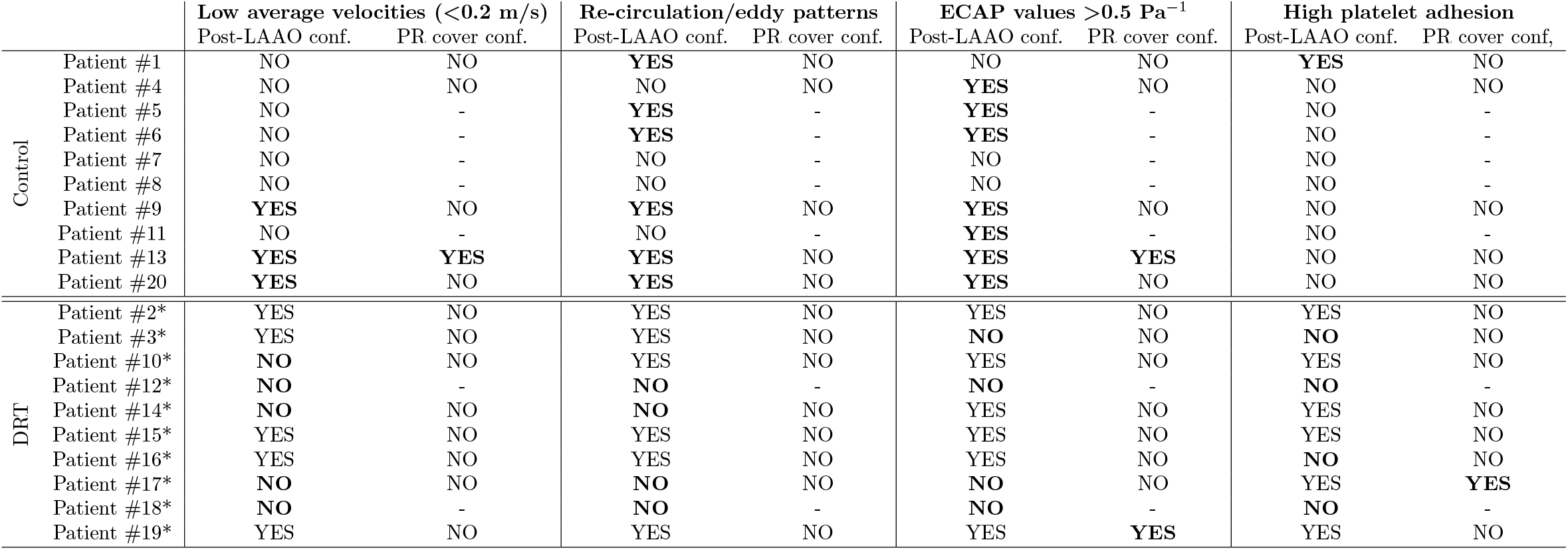
In-silico indices obtained in the 20 analysed cases with the two proposed configurations: post-left atrial appendage occlusion (LAAO) and the proposed configuration covering the PR (pulmonary ridge). Bold highlights indicate unexpected results. In the implanted post-LAAO configuration of the control group and the proposed configuration covering the pulmonary ridge, a low-risk outcome would be anticipated, where all indices should display NO. Conversely, the DRT group would be expected to exhibit the opposite. ECAP: endothelial cell activation potential * defines the patients with DRT.

## Discussion

Computational fluid dynamics simulations have the potential to aid in early diagnosis, prognosis and expedite clinical decision-making. However, to establish credibility, it is essential to personalize and verify in-silico models based on the V&V40 guidelines [47], especially in the context of health. The vast number of parameters involved in the modeling of thrombus formation in-silico presents significant challenges. Furthermore, simulating the implantation of an occluder device introduces additional variables (i.e., device type, size, location) and complexity that must be considered; thus, very few LAAO modelling contributions can be found in the literature. The main goal of this study was to develop a modeling pipeline with left atrial geometries and implanted device configurations, also allowing to modify parameters such as device type, location, and size. To our knowledge, this is the largest cohort of LAAO-based fluid simulations on patient-specific geometries. The modeling pipeline allowed to analyse the impact of the type of LAAO device (plug/pacifier) and pulmonary ridge coverage in blood flow patterns of the left atria.

The analysed cohort of patients with an implanted LAAO was balanced with 10 cases with and without DRT, respectively. In the literature, some studies have proposed in-silico hemodynamic indices such as the ECAP for DRT prediction [31, 46]. However, simulated blood flow velocities and ECAP distribution around the LAAO device did not predict DRT risk with high accuracy in the analysed cohort. Only DRT patients #2*, #15*, and #19* were clearly identified at high risk of thrombus formation after LAAO implantation due to the re-circulations at low blood flow velocities around the device, deriving into high values of ECAP, in the fluid simulations. At the other side of the spectrum, none of the in-silico descriptors could explain the DRT present in patients #12* and #18*, which had a proximal positioning in the LAA (e.g., closer to the LAA ostium) of the implanted occluder pacifier-type device. The use of generic, boundary conditions in the present study, compared to patient-specific ones in others (e.g., mitral valve velocities from echocardiography [46]) is the most plausible reason for the discrepancy in DRT prediction accuracy of fluid simulations. Moreover, DRT can eventually form in device parts such as the tip that have not been included in the generated CAD models. Additionally, no patient-specific rheological characteristics of the studied patients were available from clinical records. As a result, our fluid simulations may have underestimated the blood hypercoagulability state with a Newtonian definition, as indicated by Gonzalo et al. (2022) [25].

Despite the described simplifications of the modeling pipeline and the limitations of some in-silico hemodynamic indices, it was still useful to compare different LAAO device configurations in each individual, and analyse the impact of covering the pulmonary ridge with the LAAO device, as a preventive factor reducing the risk of DRT, as reported by Freixa et al. (2021) [16]. In our simulations, re-circulations at low blood flow velocities that may promote flow stagnation and can trigger the inflammatory process [42], were mainly found in LAAO devices positioned deeply into the LAA cavity (e.g., distal to the LAA ostium), independently of the device type. These harmful hemodynamic conditions were generally found in the upper region of the device, in agreement with the literature, which reports that 82% of DRT cases involve an uncovered left upper pulmonary ridge [16, 17]. However, depending on the individual morphological characteristics, uncovering the pulmonary ridge may not lead to DRT. For example, the LA/LAA anatomy of patients #14* and #17* (i.e., large and circular ostium, with LAA neck diameter gradually decreasing), together with the implanted LAAO positioning, led to blood flow patterns with a vortex with high flow velocities at the disk of the pacifier-type device (see Fig. 3 Patient #14* t = 0.5 s and 0.7 s), preventing the formation of re-circulations with slow blood flow. Two peri-device leaks were reported in the clinical reports (patients #2* and #7) and were maintained in the cohort, both of which involved pacifier-type occluders.

The employed discrete phase model (DPM) provided a valuable and complementary perspective in DRT risk stratification, improving the distinction between control and DRT cases. Although the threshold on platelet number and position was empirically set and its generalisation should be further tested in external databases, platelet-based in-silico indices better identified DRT patients than when using ECAP and blood flow velocities. Additionally, they also revealed a higher risk of thrombus formation for patients with uncovered pulmonary ridge, consistent with prior studies [30]. The DPM provided a clearer understanding of the device’s performance in each case, identifying susceptible areas, such as the curvature of the plug-type device, or specific cases with possible leakages. Despite the correlation between a higher number of adhered particles and cases with uncovered PR and low blood flow velocities, there were four DRT cases with particular anatomies leading to uncovered PR configurations with higher attached particle velocities: patients #14* and #17*, as described above; Patient #10*, having a vortex with high blood flow velocities in a protrusion formed at the upper part of the LAA entrance; and Patient #15*, where particles only attached between the LAAO device and the LAA wall in the covered PR configuration, in a location where only low blood velocities were present.

Pacifier-type devices demonstrated a lower risk of DRT compared to plug-types, which can be attributed to the space left by the latter between the device surface and the LAA endothelial wall, often too large to fully cover the pulmonary ridge. Even in cases with a covered pulmonary ridge, plug-type devices led to higher values of ECAP and platelet accumulation. These findings are consistent with a recent clinical study [15] that reported slightly better DRT rates in pacifier-type LAAO devices than plug-type ones. The obtained results highlight the importance of selecting the most appropriate device type and positioning for each LA morphology in relation to blood stasis. However, it should be noted that a proximal LAAO device positioning to ensure pulmonary ridge coverage may not always be feasible due to potential risks, such as improper device compression, circumflex obstruction, or insufficient endothelialization, which may increase the risk of mitral valve leaflet intersection with the pacifier disk or device embolization.

Certain limitations of the developed modeling pipeline and analysed data should be acknowledged. Variations in anticoagulant therapy and CT scan acquisition timings across the cohort may impact DRT prediction. Furthermore, the representation of LA wall movement using a spring-based dynamic mesh, based on the passive longitudinal movement observed at the mitral valve, may not fully capture the importance of 4D LA wall motion in hemodynamics and in-silico indices behavior, as highlighted by a recent sensitivity analysis on wall motion [28]. Although CT scans are commonly used for pre- and intra-LAAO planning and follow-ups, dynamic CTs providing heart motion information are not routinely included in patient protocols within the clinical routine. Echocardiography images should be explored in the future for extracting information on left atrial wall motion, as well as advanced mathematical techniques such as parallel transport [48]. Furthermore, some modeling choices in the developed pipeline such as the number of cardiac cycles, integration time, or the lack of mesh boundary layers, would not provide the most numerically robust simulations, according to recent studies [24]. However, the inclusion of more advanced modeling choices would be associated with a prohibitive computational cost if tens of patient-specific geometries are analysed. Moreover, it has already been demonstrated that having patient-specific boundary conditions is more relevant for DRT prediction than other modeling choices [46].

In this study, specific assumptions were made at the device deployment process. The web-based platform VIDAA [19] was integrated into the pipeline process, which helped in the device sizing suggestions and CAD-based device placement of the real segmented post-LAAO configuration and in the proposed configuration covering the PR to create the virtual database of simulations. Moreover, manufacturer’s recommendations and the input from an interventional cardiologist contributed to ensure a plausible device configuration, especially in the 13 proposed configurations with simulation outcomes that minimized DRT risk. However, the manual procedure employed in this study to deploy the occluder devices in the LA/LAA geometry does not account for direct interaction with the left atrial wall nor realistic deformation of the device. To achieve a more comprehensive analysis, patient-specific structural mechanical models with device-wall interaction should be investigated, similar to the ones used in the FEops HEARTguide™ platform [49], although setting up device and wall material properties is challenging. Moreover, despite preserving the main characteristics of the device shape, some simplifications were made (i.e., the hugs and catheter tip of the devices were removed, and sealing of the posterior part of the plug-type device) to reduce computational cost.

To add complexity to the discrete phase model, future research could consider integrating shear stress attachment, using particle stripping, or utilizing agent-based models in the simulations for a more realistic modeling of the thrombus formation process [50]. This would prevent particle attachment at high velocities and particle loss at high shear stress values while representing more realistic thresholds for platelet number and position related to DRT. Although simplifications are intrinsic to any modeling exercise, the integration of real post-LAAO configurations with patient-specific morphological information and representative AF boundary conditions extracted from a real patient provided valuable estimations of device configurations related to DRT risk after LAAO implantation.

## Conclusion

A computational fluid dynamics pipeline to model left atrial appendage occlusion interventions in patient-specific geometries was developed. Two types of occluder devices (i.e., pacifier versus plug shape) and positioning (i.e., coverage versus non-coverage of pulmonary ridge) were tested in 20 patient-specific left atria geometries to analyse their potential correlation with device-related thrombus formation. The obtained in-silico results confirmed that covering the pulmonary ridge with the LAAO device, when feasible, may reduce the presence of thrombogenic patterns associated with the presence of DRT. In addition, pacifier-type devices showed a better adaptability to LA/LAA morphological complexity, yielding more favorable hemodynamic results than plug-type devices, i.e., the former being associated with a reduced risk of thrombus formation. The employed modeling pipeline was not sufficiently accurate to provide a reliable prediction of DRT risk for each individual, due to the lack of patient-specific hemodynamic boundary conditions, which are required to obtain more realistic blood flow patterns. Additionally, incorporating thrombus formation models, or shear stress in the platelet adhesion models, as well as generating ground-truth data with advanced in-vitro models for model validation purposes, would further increase the credibility of the developed fluid modeling pipeline so that it can be used in the future as part of the design and regulation processes of left atrial appendage occluder devices.

## Supplementary Material

Simulated fluid flow patterns during late systole (t = 0.3 s), early diastole (t = 0.5 s), and late diastole (t = 0.7 s) in Patients #1, #2*, #3*, #4, #5, #6, #7, #8, #10*, #11, #12*, #13, #15*, #16*, #17*, #18*, #19*, #20. LAAO: left atrial appendage occlusion. * indicates patients with DRT.

## Acknowledgments

This project has received funding from the European Union’s Horizon 2020 research and innovation programme under grant agreement No 101016496 (SimCardioTest).

## A Appendix Discrete phase model effects on blood flow behaviour

**Fig A.1.**
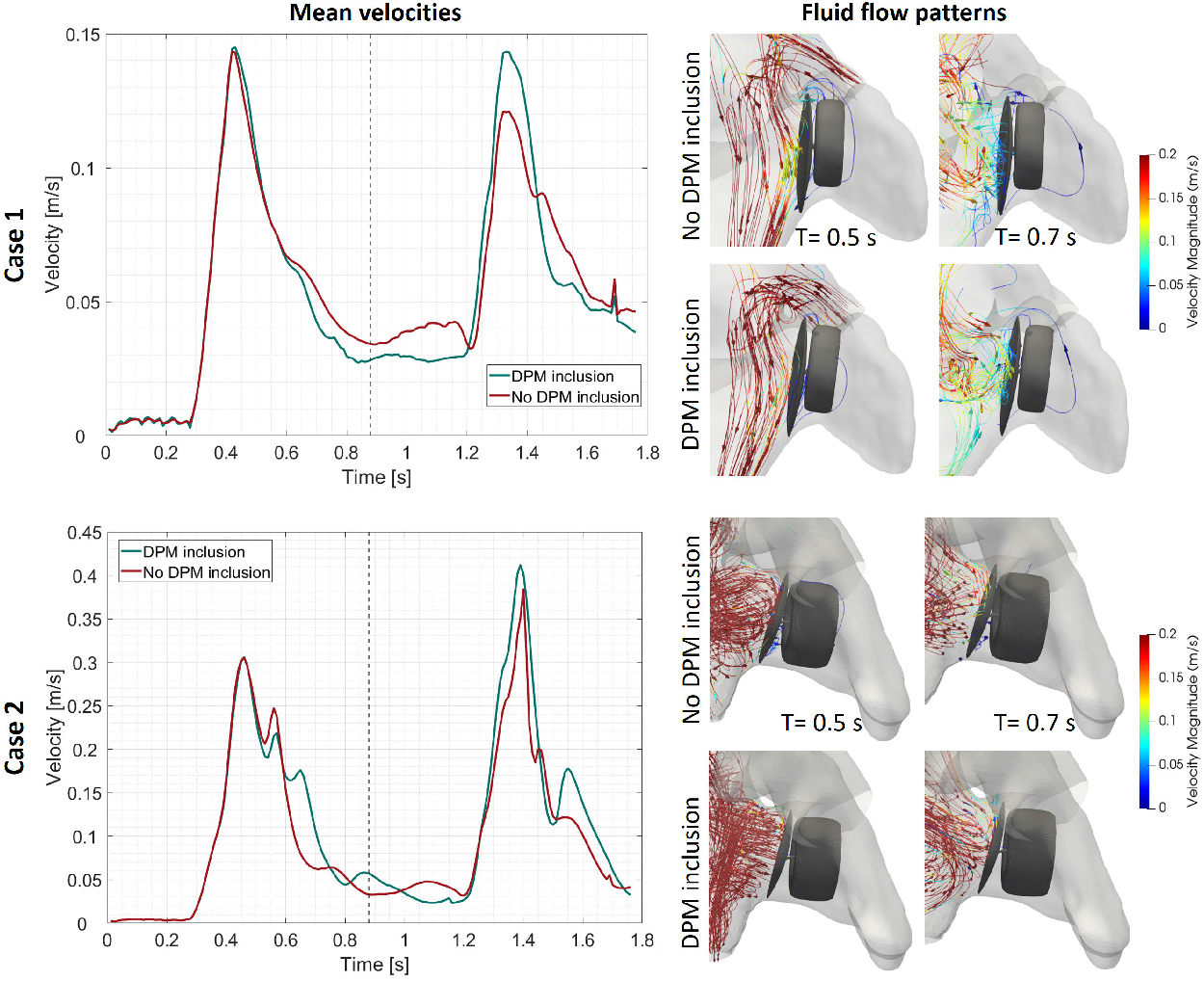
Representation of the impact of including the Discrete Phase Model (DPM) within the fluid solution domain. The left side shows a quantitative comparison of the average velocities close to the device surface within 2 cardiac cycle beats, with red indicating simulations without DPM inclusion and blue indicating simulations with DPM inclusion. The right side shows simulated blood flow patterns during early- (t = 0.5 s) and late-diastole (t = 0.7 s).

A preliminary analysis was conducted to observe any possible effects of the DPM introduction in the CFD solver on blood flow behavior. Hence, two scenarios were defined per case in two patients, introducing the DPM to interact with the LA domain fluid solution and without introducing it.

Analysing simulation outcomes in both cases, it was found that the inclusion of the DPM did not cause a noteworthy impact on the flow behavior that could alter the interpretation or estimation of DRT. Specifically, the averaged velocities at the device surface did not significantly changed throughout the cardiac beat (0.0574 *±* 0.0355 m/s with DPM inclusion and 0.0599 *±* 0.0264 m/s without DPM inclusion). However, a minor sustained velocity in the systolic phase and an increase in the diastolic peak were observed with the introduction of the DPM model (see Fig. A.1 graph). Qualitatively, similar flow patterns were observed in both patients (see Fig. A.1). At the beginning of diastole (t = 0.3 s), laminar flow with velocities higher than the 0.2 m/s threshold were predominant, except in the central part of the disk with velocities of 0.12 m/s for the no DPM inclusion scenario and 0.14 m/s in the DPM scenario. Blood flow at low velocities and with recirculations in the center of the device were observed during the end of diastole (t = 0.7 s) in both cases, slightly lower without the DPM inclusion.

**Fig A.2.**
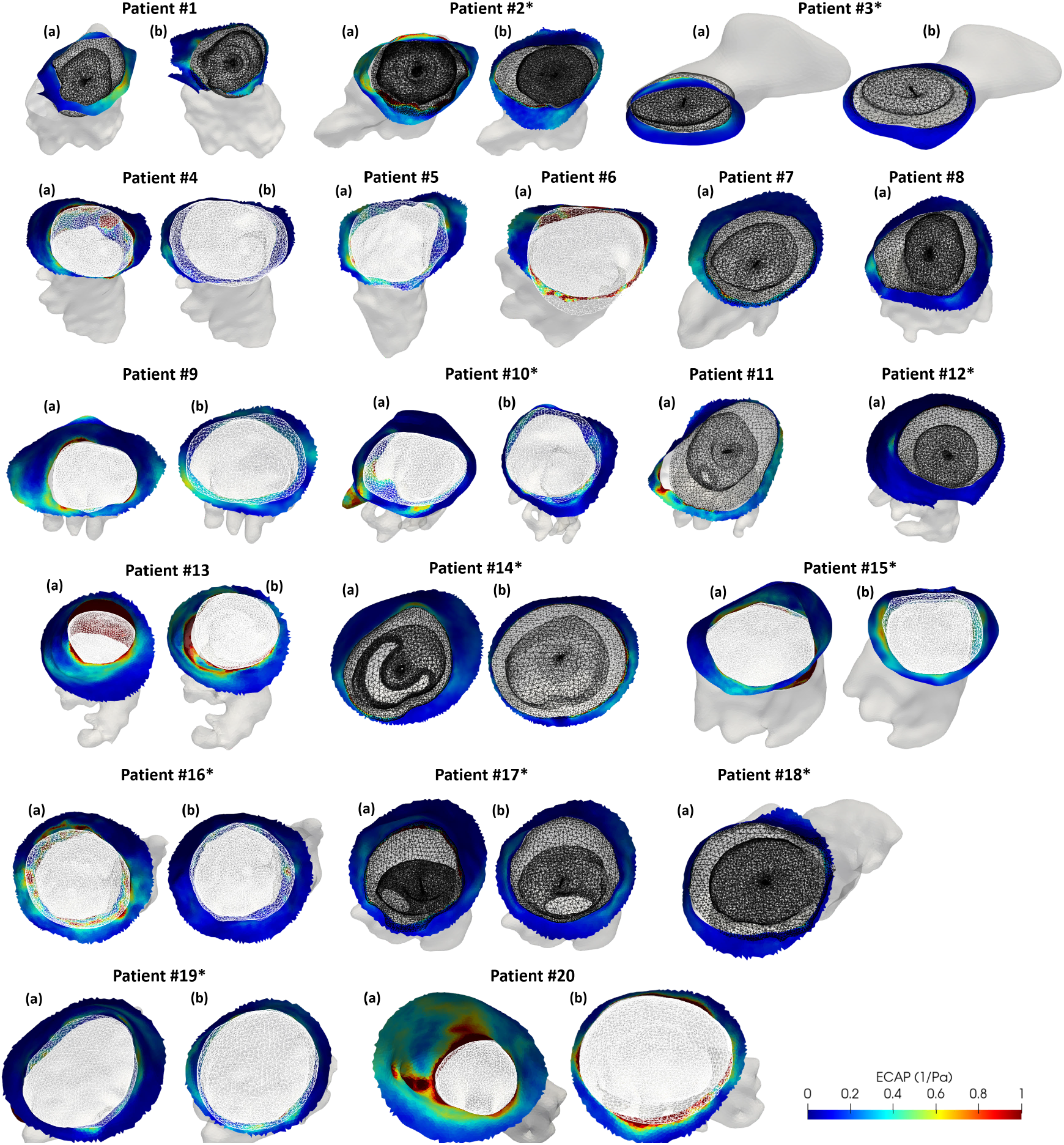
Surface representation maps of Endothelial Cell Activation Potential (ECAP) from a frontal cut of the ostium plane for the 33 analysed configurations. Device surfaces are shown in a wireframe configuration, with black representing the pacifier-type and white representing the plug-type device. Panel (a) displays the real post-Left Atrial Appendage Occlusion (LAAO) configuration, while panel (b) shows the proposed configuration covering the pulmonary ridge. * defines the patients with DRT.

**Fig A.3.**
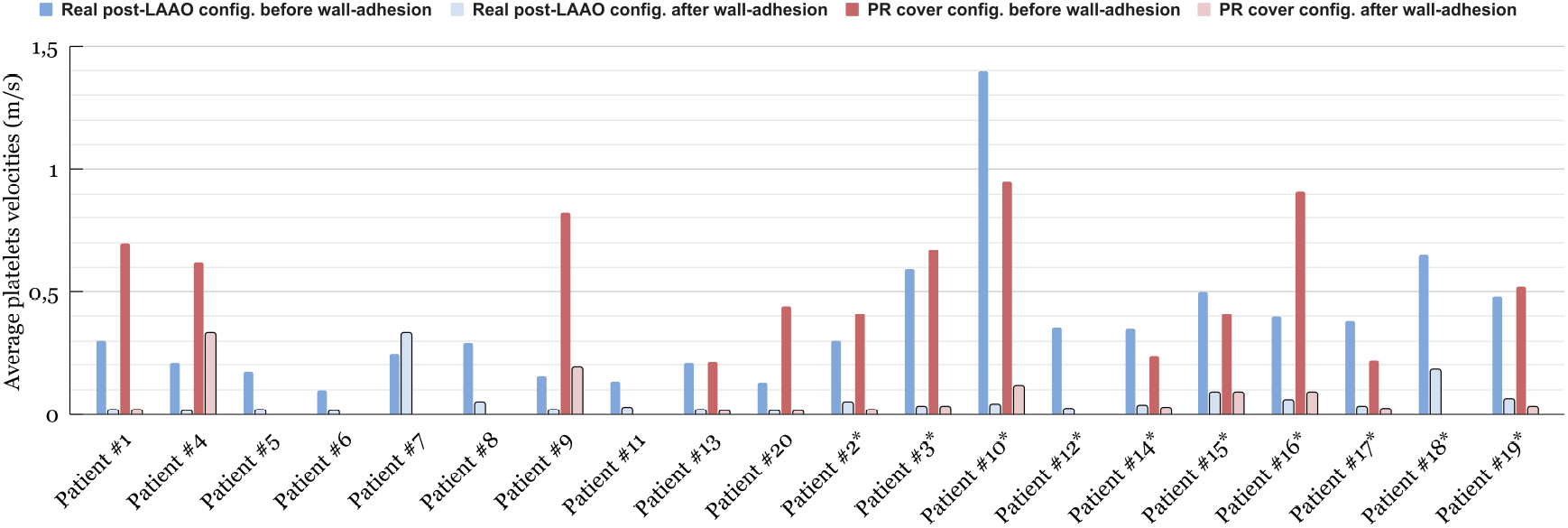
Quantification of average particle velocity before and after wall-adhesion. Blue represents the post-Left Atrial Appendage Occlusion (LAAO) configuration and, red represents the proposed configuration covering the pulmonary ridge. * denotes patients with Device-Related Thrombus (DRT).

## Footnotes

1 https://bionumbers.hms.harvard.edu/search.aspx

2 https://www.paraview.org/

